# NFAT5 participates in Inducible Nitric Oxide Synthase activation by hypoxia in MEF cells

**DOI:** 10.1101/347302

**Authors:** Yair Serman, Rodrigo A. Fuentealba, Consuelo Pasten, Jocelyn Rocco, Ben C. B. Ko, Flavio Carrión, Carlos E. Irarrázabal

**Author notes:** **Corresponding Author:** Carlos E. Irarrázabal, Ph. D; Laboratorio de Fisiología Integrativa y Molecular, Universidad de los Andes, S. Carlos Apoquindo 2200-Las Condes Santiago, Chile.

## Abstract

We previously described the protective role of NFAT5 during hypoxia, in an independent way of HIF-1α. Alternatively, inducible NO synthase (iNOS) is also induced by hypoxia. The aim of this study was to establish the NFAT5 target gene in mouse embryonic fibroblasts (MEF) cell stimulated by hypoxia. NFAT5, iNOS, NO level, aquaporin 1 (AQP1) and urea transporter 1 (UTA1) were induced by low oxygen levels in MEF cells. Additionally, NFAT5 and UTA1 were induced in reoxygenation (after 24hrs of hypoxia). NFAT5 transactivation domain (TAD) was induced during hypoxia and hypoxia/reoxygenation. Two MEF cells line independently produced for altered NFAT5 (Knockout and DBD-mutant) lost the iNOS and AQP1 induction by low oxygen. The iNOS induction was recovered in NFAT5-KO MEF cells, when recombinant NFAT5 protein expression was reconstituted, but not for NFAT5 DBD-mutant MEF cells, explained by its dominant negative effect. Finally, we found a negative feedback loop of iNOS effect over NFAT5 protein abundance. This work provides a relevant information about signaling pathway of NFAT5 during adaptive responses to oxygen depletion.

## INTRODUCTION

Hypoxia is a condition in which insufficient levels of oxygen are supplied to one or more tissues in the body. Molecular oxygen serves as the final electron acceptor in oxidative phosphorylation, and hence low oxygen supply increases the risk for generating reactive oxygen species (ROS), resulting in cell dysfunction and ultimately cell death. In this context, the nuclear factor of activated T cells 5 (NFAT5), a master regulator of the osmoprotective program, plays a protective role against hypoxia (independently of HIF-1α), but the signaling pathway is still under revision.

NFAT5 belongs to the Rel family of transcriptional activators, which includes NFκB. Cells switching to hypertonic media, NFAT5 is translocated to the nuclei [1, 2], where it binds to osmotic response elements (OREs) on target genes, increases activity of the transactivation domain (TAD) [3]. Activation of NFAT5 induces the intracellular accumulation of organic osmolytes and water transport, via induction of osmoregulatory target genes AR, UTA1, and membrane transporters AQPs, respectively [4]. In addition, NFAT5 mRNA and protein levels itself are increased by high NaCl, which confers an additional level of regulation by hypertonicity [1, 5, 6]. Besides hypertonic stimulation, NFAT5 also participates in isotonic gene expression regulation downstream of TLR4 and cytokine receptors, controlling expression of inflammatory target genes like IL-1β, IL-6, TNF-α and inducible nitric oxide synthase (iNOS) [7–10].

We have shown that NFAT5 is induced by hypoxic stimulation (0.1-2.5% O_2_), increasing its nuclear translocation and transcriptional activity. Hypoxia and hypertonicity effects on NFAT5 induction are additive, and activation of NFAT5 during hypoxia is independent of the HIF1α pathway [11]. Importantly, silencing of NFAT5 during hypoxia increased cleaved caspase-3 levels, suggesting a protective role of NFAT5 during hypoxic stimulation [11].

Endogenous nitric oxide (NO) is synthesized by three isoforms of nitric oxide synthesizing enzymes (NOS). The isoform 2 (*Nos2*), also called inducible NOS (iNOS), is induced by hypoxia. iNOS use L-arginine (Arg) to metabolize it to Lcitrulline and NO. The iNOS enzyme is calcium-independent and its expression is induced by several cytokines and immune regulators, ultimately by binding of transcription factors such as NFκB to its proximal promoter [12].

Several studies have shown that hypoxia induces the expression of iNOS, which leads to the rapid production of NO. In turn, NO combines with superoxide anion to generate peroxynitrite, a highly toxic reactive nitrogen species (RNS) that causes lipid peroxidation, nitration of protein at tyrosine residues, activation of caspase-3, apoptosis and cell death [13, 14]. Whether NFAT5 is involved in this event is currently unknown. On the other hand, induction of iNOS by lipopolysaccharide (LPS) and TLR signaling causes impairment of renal NFAT5 function due to an increase in NFAT5 S-nitrosylation, which causes a reduction of NFAT5-target gene expression in the renal medulla (ClC-K1, UTA1 and AQP2) and a consecutive sepsis-induced urinary concentration defect [15], indicating that iNOS can regulate NFAT5 activity during sepsis. Although this evidence points toward a functional reciprocity between these two signaling pathways, interaction of NFAT5 and iNOS signaling in the setting of hypoxia remains completely unexplored.

Now we investigated the relationship between NFAT5 and iNOS in MEF wild type, knockout and mutant NFAT5 during hypoxia conditions. We found that protein expression and activity of NFAT5 was induced by hypoxia in MEF cells, leading to an increase in the expression of iNOS, AQP1 and UTA1, but not of AR or BGT1. Transfection of recombinant NFAT5 in MEF-NFAT5^-/-^ cell recovered iNOS expression and partially rescued AQP1 expression during hypoxia. However, rescue was not observed in the MEF-NFAT5^Δ/Δ^ cells. A negative feedback loop of iNOS over NFAT5 was also found.

## MATERIALS AND METHODS

### Cell lines

Wild type mouse embryonic fibroblasts (MEF-NFAT5^+/+^), MEF knockout for NFAT5 (MEF-NFAT5^-/-^)[16], and NFAT5-null MEF cells expressing a mutated version of NFAT5 with skipped exons 6-7 coding for aminoacids 254-380 (MEF-NFAT5^Δ/Δ^)[17] were used.

### Cell culture treatment

Cell lines were all maintained in DMEM media (Sigma) supplemented with 2 mM L-Glutamine (HyClone), 1x Penicillin-Streptomycin (Gibco) and 10% Fetal Bovine Serum (Sigma). Pharmacological studies were done using aminoguanidine (AG, Tocris; 1mM) or L-Arg (500μM). Nitric oxide (NO) detection was performed in phenol red-free Opti-MEM1 using 5 μM of DAF-FM (Thermo). As a positive control for iNOS expression, incubation for 24 hours with 500 ng/mL LPS and 25 ng/mL IFNγ was used.

### Hypoxia and reoxygenation in cell culture

The hypoxia condition was obtained using a Heracell 150i CO_2_ incubator (Thermos) coupled to a Drager-X-am Multi-Gas Detector system set at 1% O_2_; 5% CO_2_; and 37°C [11]. A time course ranging from 4 to 48 hours was performed. The reoxygenation study was conducted after 24 hours of hypoxia with time course of reoxygenation (5% CO_2_; 21% O_2_; and 37°C) ranging from 8 to 48 hours.

### Plasmids and cell transfection

Transfection of MEF cells with Lipofectamine 3000 (Invitrogen) were done following manufacturer’s instructions. The transfection efficiency was evaluated by GFP expression in a fluorescence microscope (530 nm). Cells were fixed at room temperature for 20 minutes using PFA diluted to 4% in PBS (Sigma), followed by 4 PBS washes and a final PBS incubation until imaged. Cells were registered and counted using a fluorescence-inverted microscope (Eclipse TE2000, NIKON), and image processing and analysis were performed with NIS Elements 4.0 software and ImageJ software (NIH). Routinely, transfection efficiencies were ~50%.

### Real-time PCR (qPCR)

RNeasy mini kit (Qiagen) was used for total RNA extraction according to the manufacturer’s instructions. After verification of RNA integrity by agarose gel electrophoresis, 1 μg of RNA was reverse transcribed using a reverse transcription system (MaxyGene thermal cycler, AxyGene). The amplicons were detected by fluorescence detection by Real Time (Rotor-Gene Q, Qiagen). The results were normalized to the 18S gene.

### Luciferase assay

The binary GAL4 reporter system has been previously described [3]. To assess the activity of the NFAT5-TAD, cells were co-transfected with the GAL4 reporter (pFR-Luc) and GAL4dbd-548-1531, which contains the recombinant TAD. 24 h after transfection, cells were switched to hypoxia and luciferase activity was measured 16 h later using the Luciferase Assay System kit (Promega) and a Synergy 2 multi-detector microplate reader (Biotek).

### Cell lysate preparation and Western blot analysis of mutant and wt MEF cells

Adherent cells were lysed with cold PBS (1% TX-100), supplemented with protease inhibitor cocktail (Roche) and 1 mM PMSF (Sigma). Lysates were centrifuged at 10.000*xg*, 10 min at 4°C. Cleared lysates were determined using BCA Protein Assay kit (Pierce). The pellet was maintained at -80°C for additional analysis. Equal amounts of soluble proteins were separated by SDS-PAGE and transferred to a PVDF or nitrocellulose membrane. Membranes were blocked with Odyssey blocking buffer: PBS (1: 4), and overnight incubated at 4°C with primary antibodies in Odyssey blocking buffer. Next day, membranes were incubated 2 hours with fluorophore-conjugated secondary antibody (Invitrogen). Proteins were detected by Infra-Red fluorescence (IR) using the Odyssey CLx system (Li-Cor). The intensities of the resulting bands were quantitated by infrared fluorescence signal (Image Studio Lite version 5.25; Li-Cor). The antibodies used were rabbit anti-NFAT5 (Thermos), mouse anti-NFAT5 (SantaCruz), mouse anti-V5 (Invitrogen), mouse anti-iNOS (BD Transduction Laboratories), rabbit anti-AQP1 (Abcam), rabbit anti-UTA-1 (Abcam), rabbit anti-3-NT (Millipore), or rabbit anti-β-actin (Sigma) antibodies.

### Statistical Analysis

Data are expressed as average + SEM. Values from different groups were assessed with 1 way Anova test for multiple comparisons with a post-hoc Tukey. The significance level was *\*, p*≤ 0.05, *\*\*, p*≤ 0.01; or *\*\*\*, p*≤ 0.001.

## RESULTS

### Hypoxia induces a time-dependent increase of NFAT5 and iNOS expression in MEF cells

A time course of hypoxia on NFAT5 and iNOS expression was measured by RT-qPCR and Western blot (Figure 1). Levels of NFAT5 mRNA increased 3.8 times at 4 hours of hypoxia (Figure 1A), returning it to baseline values at 16 hours after hypoxia induction. During basal conditions (21% O_2_), detectable levels of NFAT5 protein were found in MEF cells (Figure 1B). Compared to normoxia, hypoxia (1% O_2_) produced a significant increase of NFAT5 protein levels (40%) 4 hours of hypoxia, then the protein levels returned to baseline values and a second peak of NFTA5 expression was observed (24 and 48 hours after hypoxia; 30%) (Figure 1B).

**Figure 1.**
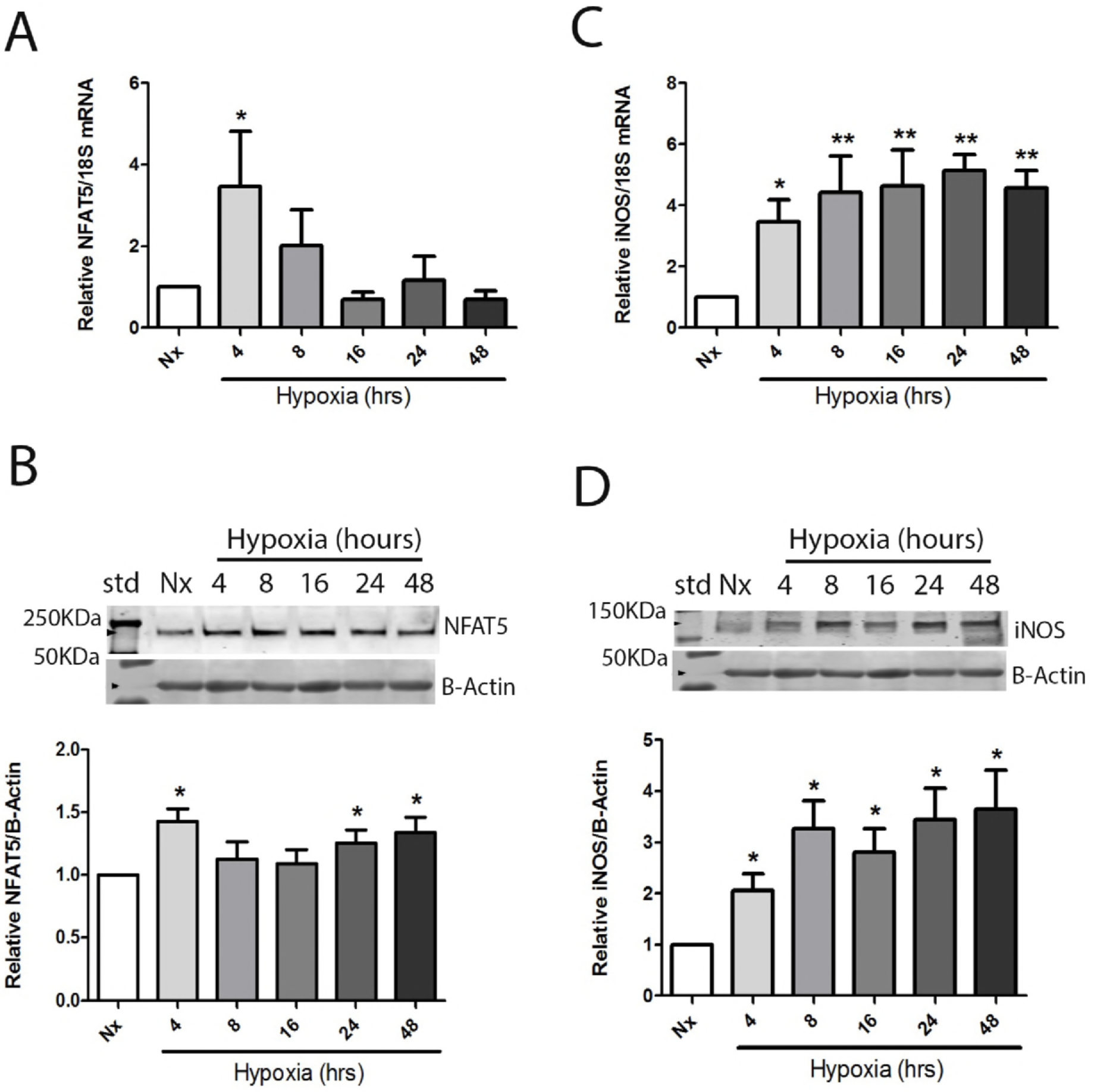
Hypoxia increases mRNA and protein levels of NFAT5 and iNOS in MEF cells. Wild type MEF cells were incubated in normoxia conditions (Nx, 21% O_2_) or time points of hypoxia (4-48 h, 1%O_2_). Relative mRNA or protein levels for NFAT5 and iNOS were determined by RT-qPCR (rotorgene, Qiagen) or quantitative Western blot (LicoR system), respectively. *(A)* Relative NFAT5 mRNA. *(B)* Relative NFAT5 protein. *(C)* Relative mRNA levels of iNOS. *(D)* Relative protein levels of iNOS. Mean ± SEM; *n* = 7. *\*, p* ≤ 0.05; *\*\*, p* ≤ 0.01 compared to normoxia.

During basal conditions (21% O_2_) iNOS expression is very low in MEF cells. The iNOS mRNA increased 3.9 times at 4 hours of hypoxia and it was sustained in time points analyzed (Figure 1C). At 4 hrs of hypoxia, iNOS protein levels were significantly induced, reaching a plateau of expression from 8 hours of hypoxia (3- fold over normoxia levels) (Figure 1D). According with these results hypoxia induced NFAT5 and iNOS during 4-48 hours period of hypoxia. We established 24 hours of hypoxia to study the hypoxia/reoxygenation experiments in MEF cells.

### Time course effect of reoxygenation on NFAT5 and iNOS expression in MEF cells

After 24 h of hypoxia, a time course of reoxygenation was conducted. NFAT5 mRNA levels decreased rapidly after switching to 21% O_2_ (Figure 2A). However, levels of NFTA5 protein was transient induced during reoxygenation when compared to hypoxia. NFAT5 increased significantly from 8 h reoxygenation, peaking at 16 h, and then returning to the levels observed during hypoxia condition at 24-48 h (Figure 2B). The iNOS mRNA levels experimented a gradual decrease reaching a baseline level 48 hours after reoxygenation (Figure 2C). On the other hand, the iNOS protein expression levels fall rapidly during reoxygenation, but interestingly remained significantly above the baseline level (1.8 times) (Figure 2D).

**Figure 2.**
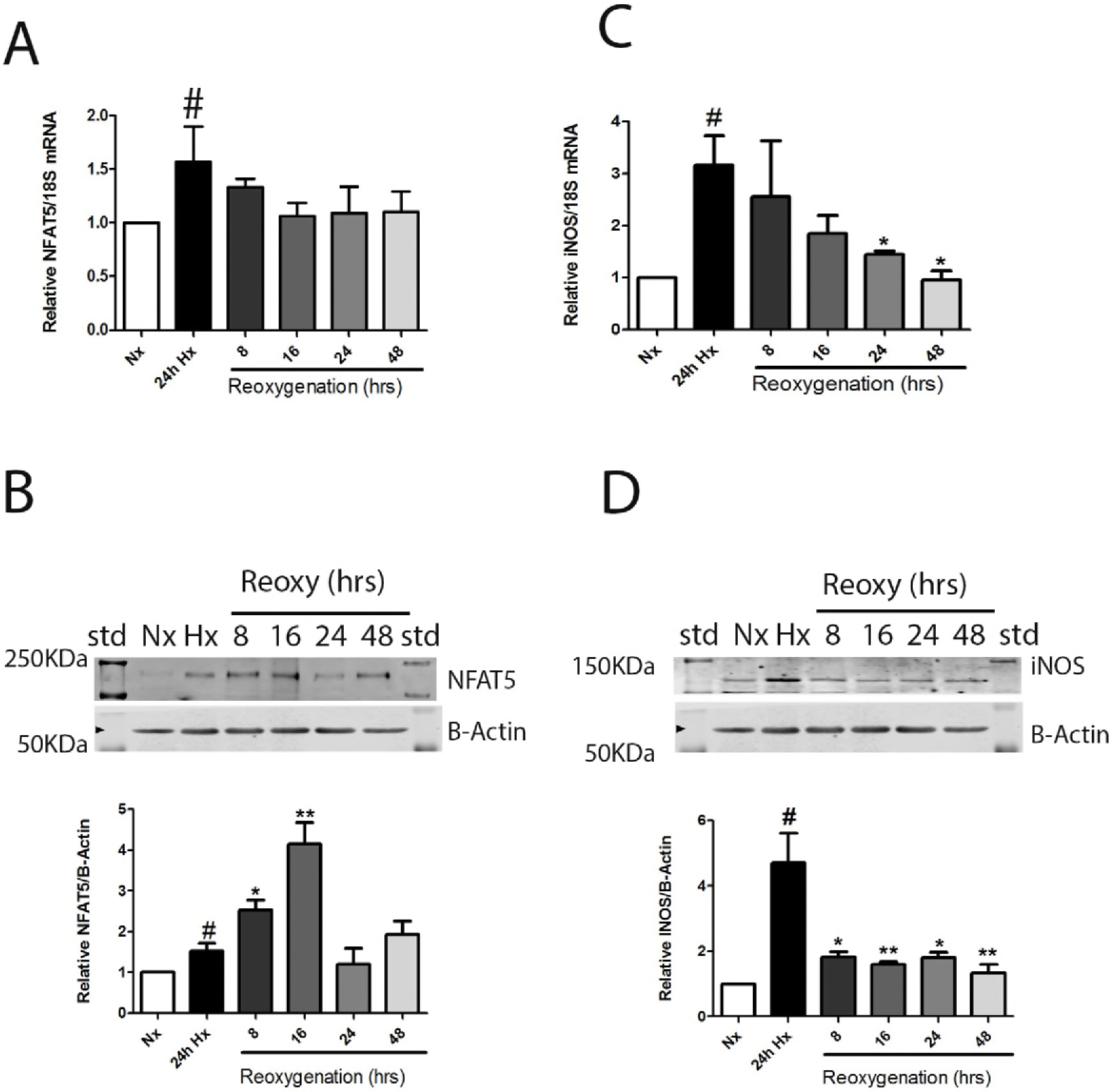
Induction of NFAT5 and iNOS during hypoxia/reoxygenation. MEF wild type cells were incubated in normoxia (Nx, 21% O_2_), 24 h of hypoxia (Hx, 1% O_2_), or time points of reoxygenation (8-48 h, 21%O2) after an initial 24 h hypoxia 1% O_2_. *(A)* Relative mRNA expression of NFAT5. *(B)* Relative protein expression of NFAT5. *(C)* Relative mRNA levels of iNOS. *(D)* Relative protein expression of iNOS. Mean ± SEM; *n* = 7. *#, p* ≤ 0.05 compared with normoxia; *\*, p* ≤ 0.01 compared with hypoxia.

### NFAT5 transcriptional activity was induced by hypoxia and hypoxia/reoxygenation

Previously, we demonstrated that NFAT5 knockdown decreased HRE-Luciferase activity [11]. Now our results established that our transfection protocol had a 50% of transfection efficiency (GFP expression). The results showed that empty vector (pFRLuc co-transfected with pFA) did not produce a detectable luciferase activity. However, when cells were transfected with NFAT5-TAD a basal activity was observed in normoxia condition (20 URL, Figure 3). However, MEF cells exposed to hypoxia (24h), experimented higher transactivation activity than normoxia (52 URLs). Hypoxia/reoxygenation (24/16h) increased transactivation activity relative to basal condition (similar levels to observed in hypoxia). The positive control of transactivation activity reached an average of 170 URLs (16h hypertonicity) (Figure 3).

**Figure 3.**
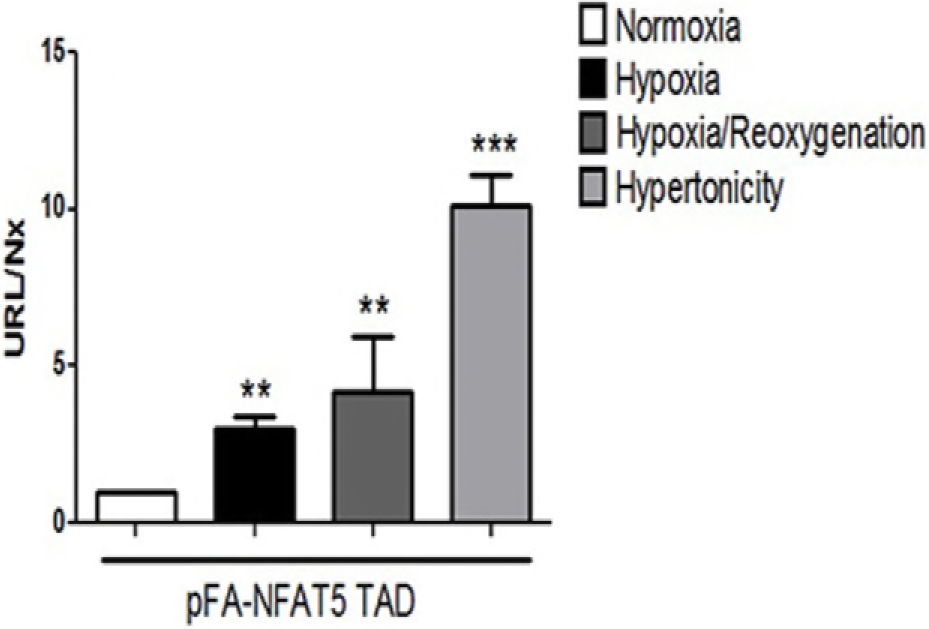
Hypoxia and hypoxia/reoxygenation do increase NFAT5 transcriptional activity. MEF cells co-transfected with pFR-LUC plus pFA-NFAT5-TAD. Graph bars represent normalized URLs for the different conditions tested. Mean ± SEM; *n =* 6. *\*\*, p* ≤ 0.01; *\*\*\*, p* ≤ 0.001 compared to normoxia condition.

### NFAT5 target genes activated by hypoxia and hypoxia/reoxygenation

We observed that protein levels of AQP-1 and UTA-1 were significantly induced 24 hours of hypoxia (Figure 4A, 4B), but neither AR nor BGT1 changed its protein levels (Figure 4C and 4D). The mRNA levels of AQP1 and UTA1 were increased by 24 h of hypoxia too (Figure 4E and 4F). The proteins levels of AQP1 returned to normoxia levels during reoxygenation (Figure 5A). UTA1 protein abundance was significantly induced after 16 hours of reoxygenation compared with hypoxia (Figures 5B). AR and BGT1 did not experimented changes in its expression during reoxygenation step (Figure 5C y 5D). The AQP1 mRNA level after 8 hrs of reoxygenation were high and equivalent to those detected during hypoxia, but then they fall gradually toward 48 hours (Figure 5E). UTA1 mRNA levels decreased quickly, returning to normoxia levels as soon as 8 hours after reoxygenation (Figure 5F). These results showed that in MEF cells hypoxia produced up-regulation of iNOS, AQP1 and UTA1, after NFAT5 activation, but not during reoxygenation process, except for UTA-1.

**Figure 4.**
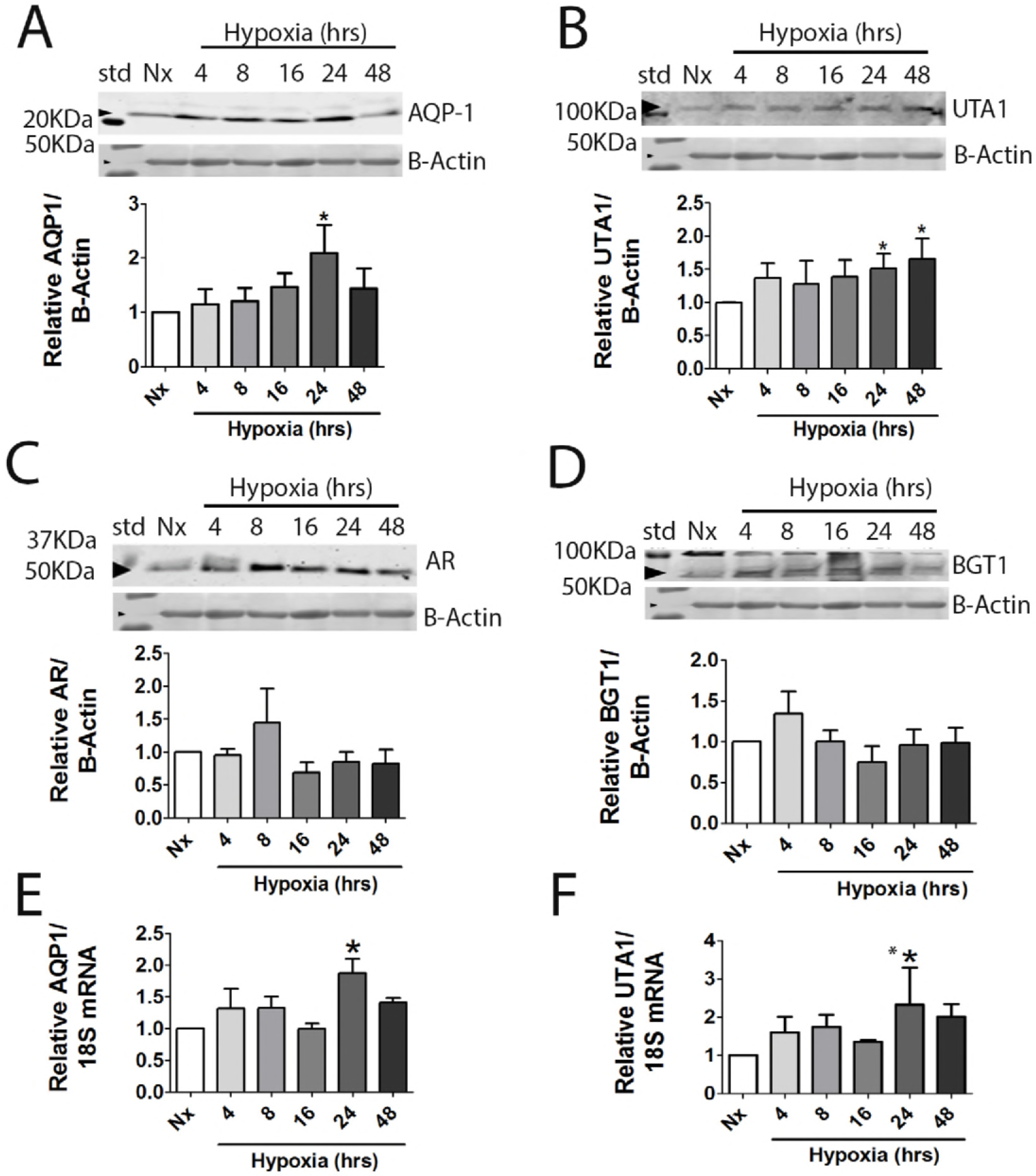
Hypoxia stimulates mRNA and protein levels of potential NFAT5-target genes. MEF wild type cells were incubated in normoxia conditions (Nx, 21%O_2_) or for different time points of hypoxia stimulation (4-48 h, 1%O_2_). *(A)* Relative AQP1 protein expression. *(B)* Relative UTA1 protein expression. *(C)* Relative AR protein expression. (D) Relative BGT1 protein expression. *(E)* Relative mRNA expression of AQP1. *(F)* Relative mRNA expression of UTA1. Mean ± SEM; *n =* 7 *\*\*, p* ≤ 0.01 *\*\*\*, p* ≤ 0.001 compared to normoxia condition.

**Figure 5.**
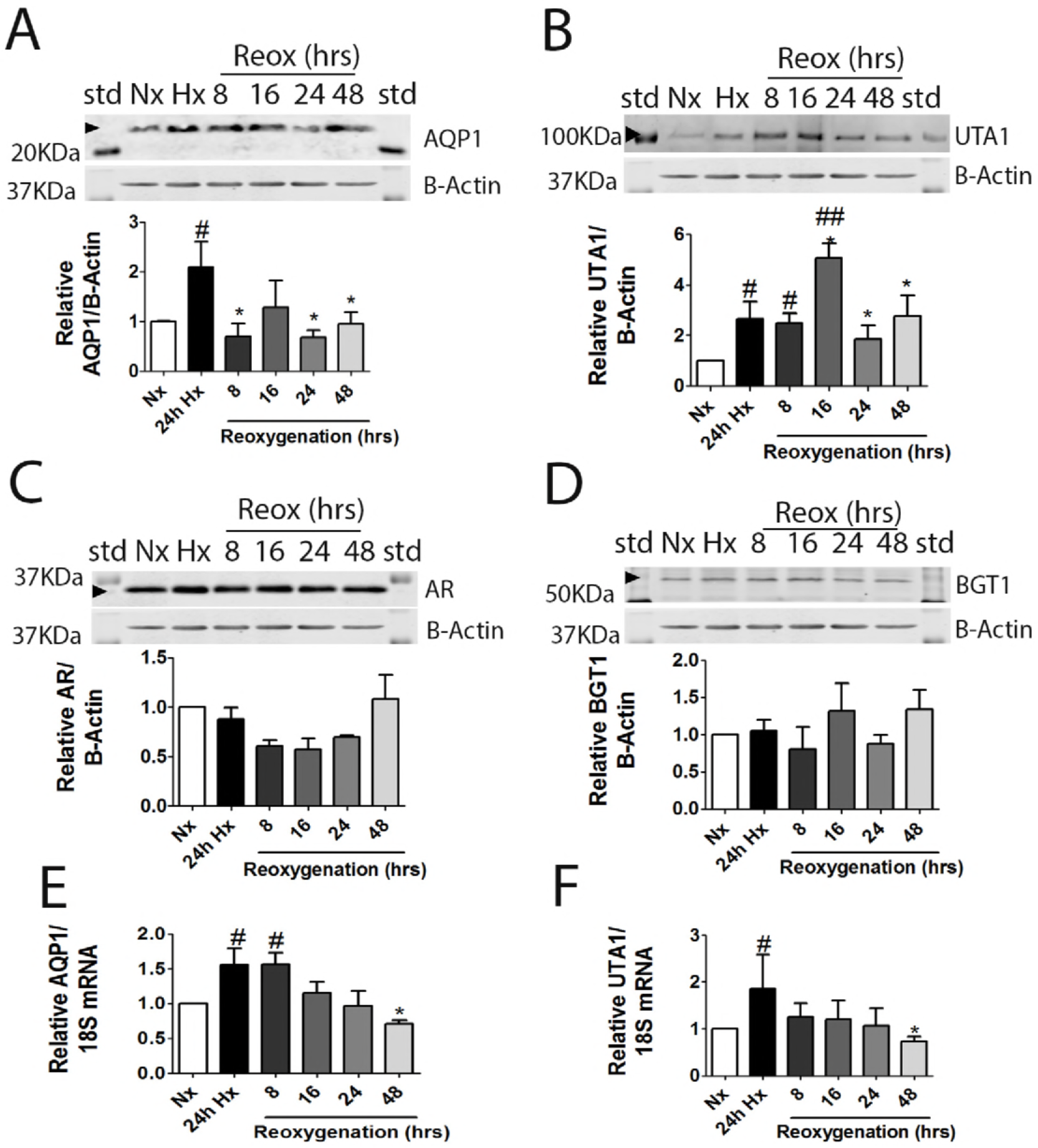
Hypoxia/reoxygenation induce potential NFAT5-target genes. MEF wild type cells were incubated in normoxia (Nx, 21% O_2_), 24 h of hypoxia (1% O_2_), or during a time course of reoxygenation (8-48 h, 21%O_2_). *(A)* AQP1 protein. *(B)* UTA1 protein. *(C)* AR protein. *(D)* BGT1 protein. *(E)* AQP1 mRNA. *(F)* UTA1 mRNA. Mean ± SEM; *n =* 7; *#, p* ≤ 0.05 compared with normoxia; *\*, p* ≤ 0.01 compared with hypoxia.

### NFAT5 target genes regulation under hypoxia conditions in MEF-NFAT5^-/-^ cells and pharmacological NO manipulation

To understand the role of NFAT5 on iNOS, AQP-1 and UTA1 induction by hypoxia, we use NFAT5 knockout cells (MEF-NFAT5^-/-^). We confirm that NFAT5-KO cells do not express NFAT5 protein (Figure 6A). The induction of iNOS and AQP-1 by hypoxia was significant eliminated in MEF-NFAT5^-/-^ cells. These results suggest the participation of NFAT5 on iNOS and AQP-1 response to hypoxia in MEF cell (Figure 6B).

**Figure 6.**
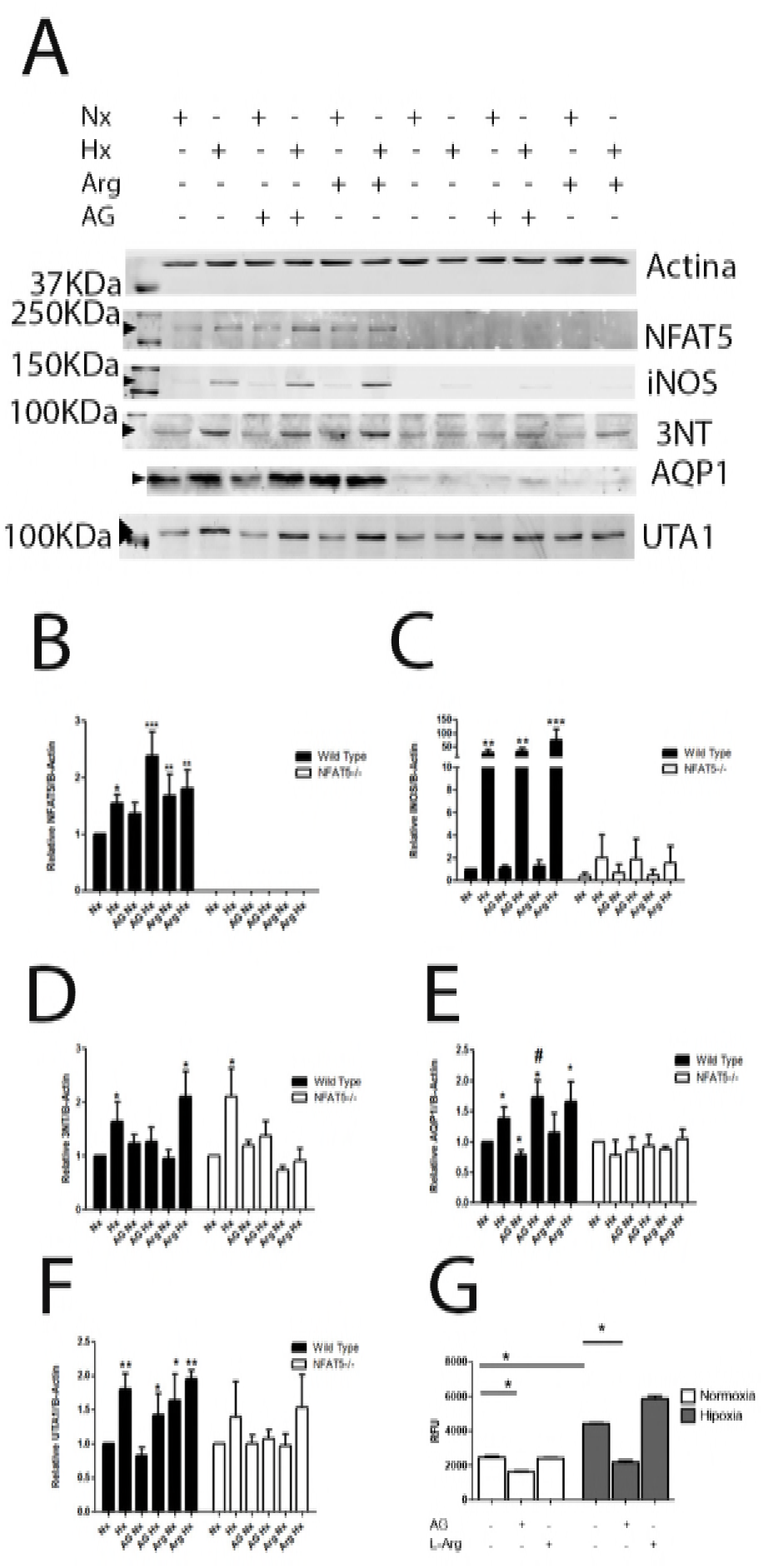
NFAT5 is required for hypoxia-stimulated increase of iNOS and AQP1 expression. MEF wild type and MEF-NFAT5^-/-^ cells were incubated in normoxia (line 1; Nx, 21% of O_2_) or hypoxia (line 2, 1% of O_2_) condition. Cells were pre-incubated for 1 h with AG (1 mM) before normoxia (line 3) or hypoxia (line 4) conditions. L-Arg (500 μM) before normoxia (line 5) or hypoxia (line 6) conditions. Graphics are normalized by normoxia condition for each cell type. *(A)* Relative protein levels were determinated by quantitative Western blot *(B)* NFAT5. *(C)* iNOS. *(D)* AQP1. *(E)* UTA1. *(F)* 3-NT, in a representative selected area. Mean ± SEM; *n =* 4, *\*, p* ≤ 0.05, *\*\*, p* ≤ 0.01, *\*\*\*, p* ≤ 0.001; relative to normoxia. *#, p* ≤ 0.05 relative to hypoxia.

We tested the iNOS activity during hypoxia, measuring Nitric Oxide (NO) levels using a NO-sensitive probe. Also, we measurement the protein nitrotyrosilation levels (a product of NO levels) using a 3-NT antibody. The production of NO and 3-NT was induced by hypoxia. Interestingly, the 3-NT levels in MEF-NFAT5^-/-^ cells were lower than in MEF-NFAT5^+/+^ cells. The preincubation with L-Arg increased the NO and 3-NT levels during hypoxia in MEF-NFAT5^+/+^ cells, but not in KO cells (Figure 6E). These results suggest that NO production in hypoxia strongly depends on NFAT5 (Figure 6B).

The iNOS expression increased 30-fold in MEF-NFAT5^+/+^ under hypoxic conditions (Figure 6C). The treatment with AG (iNOS inhibitor) did not affect the expression of iNOS in MEF cells. The AG inhibited the NO and 3-NT levels observed during hypoxia. Interestingly, during hypoxia AG increased significantly the NFAT5 protein abundance in wild type cells (Figure 6B), but not L-Arg (Figure 6B). These data suggest a interplay between Inducible Nitric Oxide Synthase and NFAT5 in hypoxic condition, as it has been described for osmotic regulation [18].

As it was mentioned above, the AQP1 abundance in MEF-NFAT5^-/-^ cells were undetectable in all conditions tested (Figure 6F), indicating that basal and hypoxia-induced AQP1 expression was NFAT5-dependent. In wild type cells, the iNOS inhibitor (AG), increased significantly the AQP1 abundance in hypoxia (72% of AQP1), showed that AG treatment concomitantly improved the NFAT5 and AQP1 expression.

Regarding UTA1 regulation, we observed that UTA1 induction by hypoxia in wild type cells was partially lost with AG (Figure 6G). On the other hand, the L-Arg treatment induced UTA1 abundance in normoxia, but did not modified their levels in hypoxia condition (Figure 6G). Interestingly, MEF-NFAT5^-/-^ cells lost the induction of UTA1 by hypoxia compared to wild type cells, regardless of pharmacological NO manipulation (Figure 6G).

### Full length human NFAT5 rescues hypoxia-induced effects on NFAT5 target genes in MEF-NFAT5^-/-^ cells, but not in MEF-NFAT5^Δ/Δ^ cells

The above results clearly suggest that the hypoxia-induced increase of AQP1 and iNOS levels were dependent on NFAT5 activity in MEF cells. To further confirm this, we used MEF cells expressing a mutated version of NFAT5 (MEF-NFAT5^Δ/Δ^). This NFAT5 mutant lacks residues 254-340 of NFAT5 and loses its ability to DNA binding [17] and enhanceosome formation [10]. In contrast to wild-type cells, the MEF-NFAT5^Δ/Δ^ cells lose iNOS induction elicited either by hypoxia or hypertonicity stimulation (Figure 7). Notably, the expression of AQP1 was also lost in all the conditions tested (normoxia, hypoxia and hypertonicity) in MEF-NFAT5^Δ/Δ^ cells. This result strongly suggests that DNA binding to cognate TonE elements by NFAT5 and NFAT5-dependent enhanceosome formation in the in *Aqp1* and *Nos2* promoter are also important during hypoxia stimulation. Collectively, our results demonstrate that the induction of AQP1 and iNOS by hypoxia stimulation were NFAT5-dependent.

**Figure 7.**
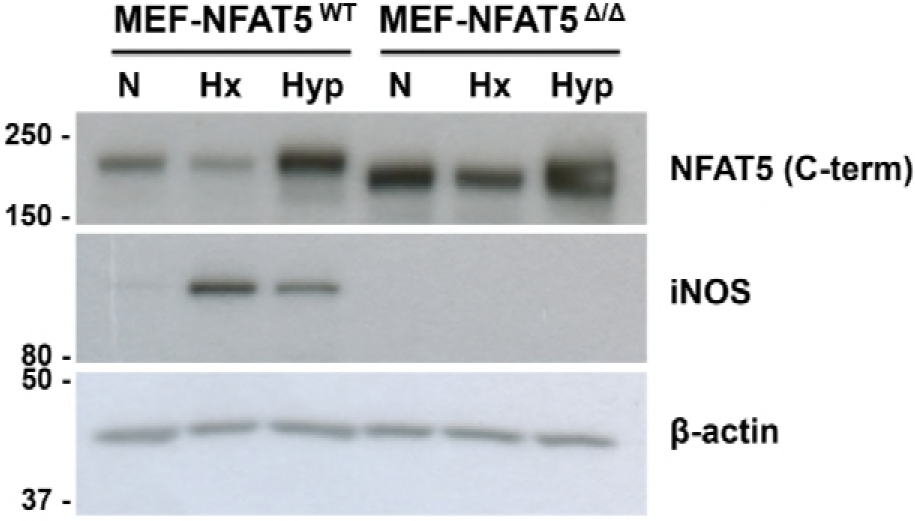
Mutant NFAT5 blunts iNOS responsiveness to hypoxia and hypertonicity. Wild type MEF cells (MEF-NFAT5^wt^) and NFAT5-null cells (MEF-NFAT5^Δ/Δ^) were subjected to Normoxia (N), Hypoxia (Hx, 1% O_2_) or Hypertonic (Hyp, 100 mM NaCl) conditions for 24 h. Images are representative of four independent experiments.

**Figure 8.**
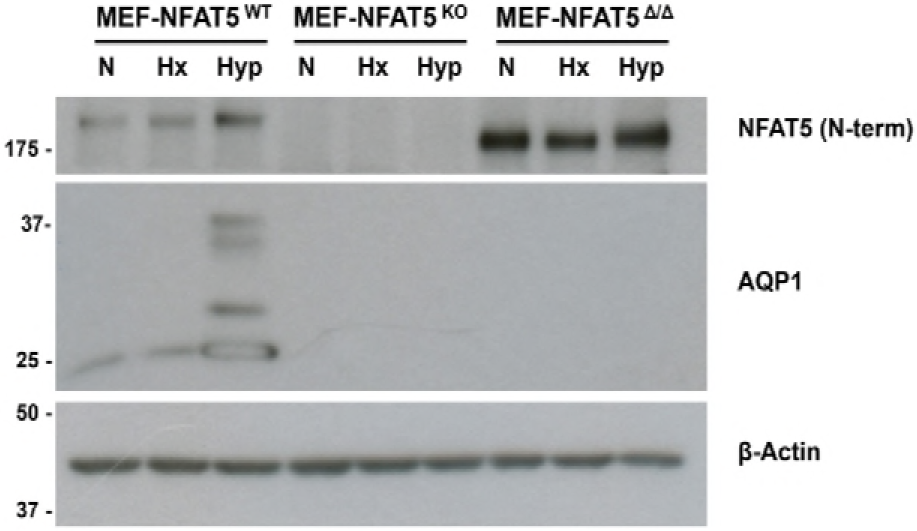
NFAT5 Knockout and Null cells are unresponsive to hypoxia- and hypertonicity-stimulated AQP1 expression. Wild type MEF cells (MEFNFAT5^wt^), NFAT5 KO (MEF-NFAT5^wt^), or NFAT5-null (MEF-NFAT5^Δ/Δ^) were subjected to Normoxia (N), Hypoxia (Hx, 1% O_2_) or Hypertonic (Hyp, 100 mM NaCl) conditions for 24 h. Images are representative of four independent experiments.

The rescue experiments showed, in MEF-NFAT5^-/-^ cells, that reconstitution of recombinant NFAT5 recovered the iNOS induction by hypoxia (Figure 9A). Levels of iNOS significantly increased upon NFAT5 re-expression, reaching twofold levels during hypoxia (Figure 9B). Noteworthy, rescue experiment did not work in MEFNFAT5^Δ/Δ^ despite an even higher expression of the recombinant full-length protein compared to levels achieved in knockout cells (Figure 9A). This striking observation suggests that iNOS expression by hypoxia depends on intact NFAT5 heterodimers, and that background NFAT5^Δ/Δ^ expression might effectively serve a dominant negative role now for hypoxia-mediated iNOS induction in MEF-NFAT5^Δ/Δ^ cells, as previously demonstrated for NFAT5-mediated isotonic gene expression regulation [10]. We also observed rescue AQP1, but in less degree than iNOS (Figure 9A).

**Figure 9.**
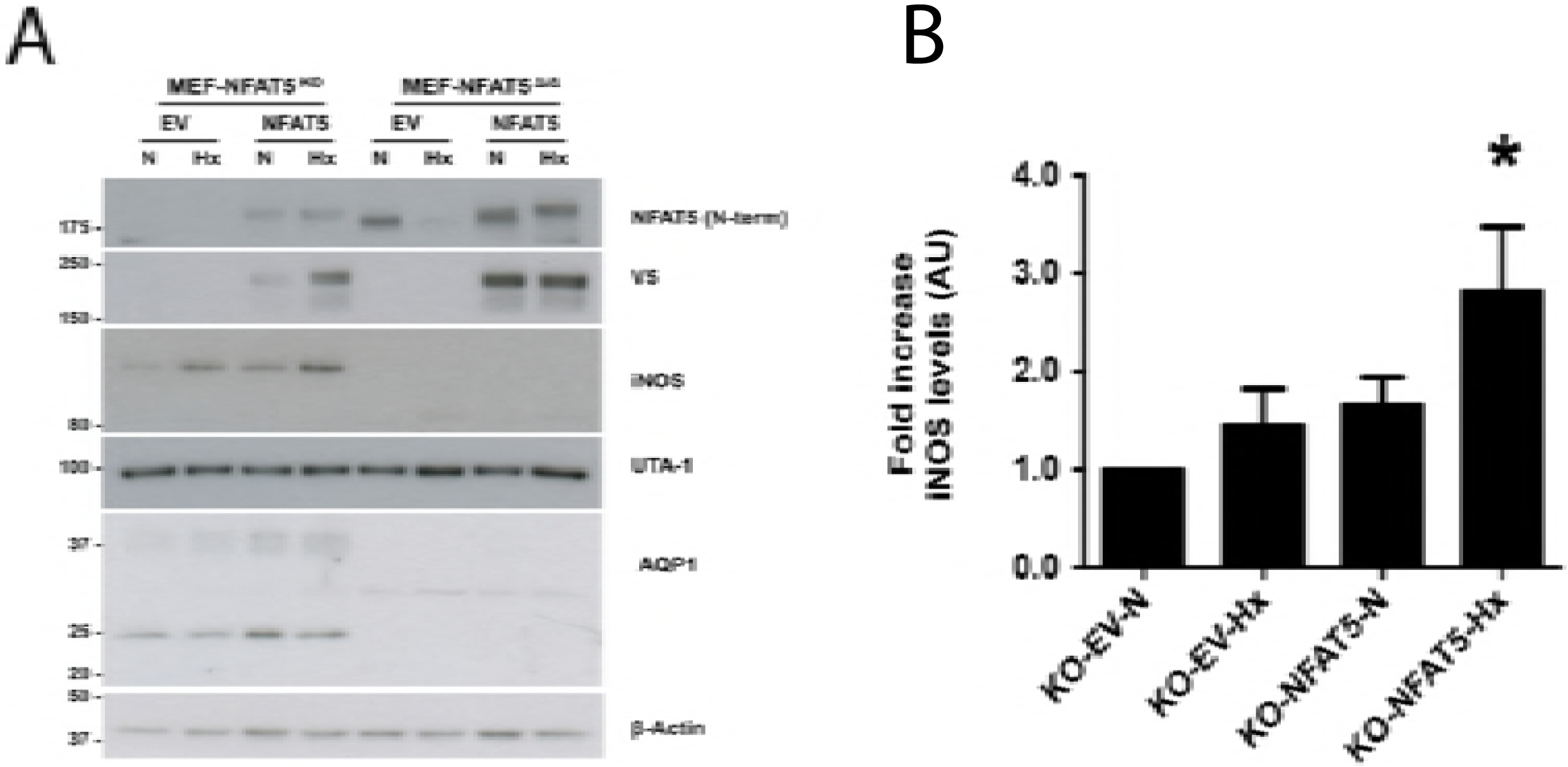
Restoration of NFAT5 expression in NFAT5 knockout cells, but not in NFAT5 null cells, induces a partial rescue of iNOS and AQP1 expression during hypoxia. A, NFAT5 knockout (MEF-NFAT5^KO^) and NFAT5-null (MEFNFAT5^Δ/Δ^) cells were transfected with recombinant NFAT5 or corresponding empty vector (EV). Cells were recovered for 16 h and then subjected to Normoxia (N) or Hypoxia (Hx, 1% O_2_) conditions for 24 h. *(A)* A representative Western blot analysis showing NFAT5 (SantaCruz anti-NFAT5 or anti-V5 antibody), iNOS, UTA1, AQP1 and β-Actin detection. *(B)* Densitometric analysis for iNOS detection from Western blots obtained from 5 independent experiments. Mean ± SEM; *\*, p* ≤ 0.05 respect of normoxia with empty vector (KO-E).

## DISCUSION

Besides a master regulator for osmoadaptative responses to hypertonic stress [4], mounting evidence supports now a leading role of NFAT5 as an essential factor for organ development [16, 19], as an integral component of the immune response [7, 17], a systemic surveillance factor for blood pressure control [20, 21], a key regulator of the urine concentration mechanism for renal function [22], and as a novel HIF1α-independent protective factor against hypoxia [11]. Therefore, NFAT5 can be considered as a pivotal transcription factor and integrator of multiple adverse stimulus for stress adaptation. Another molecule highly studied during hypoxia is NO, which mediates positive or adverse effects [23, 24]. Of the three isoforms that produce NO, iNOS enzyme is independent of intracellular calcium concentration and it has been described that it exerts a negative regulatory role for NFAT5-mediated signaling in the renal medulla during sepsis [15]. There are no studies to clarify NFAT5 and iNOS relationship in the setting of hypoxia and hypoxia/reoxygenation.

In MEF cells we found that NFAT5 and iNOS were induced shortly after hypoxia, starting 4 hours after stimulation. In the hypoxia/reoxigenation condition we found that NFAT5 presented a further increase although NFAT5 mRNA levels had returned to basal levels, suggesting protein stabilization mechanisms. However, the transactivation activity of NFAT5 was also increased in hypoxia and hypoxia/reoxygenation, confirming our previous results in HEK293 and rat primary IMCD cells [11]. Unexpectedly, reoxygenation reduced the expression of iNOS compared to hypoxia levels, but still remained higher than basal levels (1.8 times over normoxia condition). We found now that in hypoxia, AQP1 and UTA1 protein expression was increased in MEF cells. However, neither AR nor BGT1 levels changed upon stimulation, suggesting that hypertonic target genes are differentially regulated by NFAT5 during hypoxia, either via a signaling pathway independent of ORE, or being negatively affected by chromatin context during hypoxia. Regardless mechanism, our results are in agreement with previous results showing that AR is not induced by hypoxia [25].

Interestingly, the NFAT5 KO (MEF-NFAT5^-/-^) and null (MEF-NFAT5^Δ/Δ^) MEF cells did not expresses AQP1 or iNOS during hypoxia stimulation. These two cell types were produced independently by two different groups, emphasizing our findings that NFAT5 is controlling the gene expression of these two proteins during hypoxia. We investigated this finding deeper and we explored the reconstitution of AQP1 and iNOS expression in the NFAT5-deficient cells line by restoring expression of NFAT5. In KO MEF cells AQP1 and iNOS were recovered with NFAT5 reconstitution. However, the null (MEF-NFAT5^Δ/Δ^) MEF cells did not express AQP1 or iNOS even when recombinant NFAT5 was reconstituted, confirming previous report describing NFAT5 mutant as a dominant negative. Interestingly, this important result suggests that during hypoxia, iNOS and AQP1 gene expression are similarly regulated. Whether AQP1 is also stimulated via enhanceosome formation during hypoxia as for the *Nos2* gene during isotonic immune regulation [10], is currently unknown.

Our results showed that iNOS activity has a negative role on NFAT5 expression in response to hypoxia, which might be explained by a post-translational modification that increased its degradation, as it was previously described by NO-dependent modification upon LPS treatment [18]. During hypoxia, the inhibition of iNOS activity (AG), significantly increased the expression of NFAT5 and AQP1. The AG has been described as a selective inducible nitric oxide synthase inhibitor [26]. However, other multiple biological effects have been postulated, including prevention of advanced glycation end products (AGEP) formation [27], and inhibition of diamino oxidase [28]. We are currently exploring whether the effect of AG on NFTA5 and AQP1 could be related to the other biological activities of AG during hypoxia.

The expression of UTA1 at 24 hours after induction of hypoxia could be explained because hypoxia also induces arginase 2, which produces ornithine plus urea. Arginase-2 is induced in hypoxia and constitutes a protective mechanism for the deleterious effects of iNOS activity [29]. We propose that UTA1 upregulation was a secondary response to urea overproduction by hypoxia [30]. In MEF-NFAT5^-/-^ cells UTA1 is expressed in basal form, but hypoxia-UTA1 induction by was lost, suggesting that NFAT5 participates in the induction of UTA1 during hypoxia as well. We are exploring this hypothesis.

In conclusion, these results confirm that NFAT5 is activated by hypoxia and hypoxia/reoxygenation. During hypoxia NFAT5 control the expression of AQP1 and iNOS. During hypoxia/reoxygenation, NFAT5 might control UTA1 gene expression. Additionally, increased production of NO by iNOS inhibited the expression of NFAT5 during hypoxia. Therefore, there is a bidirectional communication between NFAT5 and iNOS signaling during cell adaptation to decreased oxygen supply. These results improve the understanding for future pharmacological interventions during ischemic pathology.

## Contributions

The study was conceived and designed by CEI. YS performed most experiments (including Western blot, qRT-PCR, luciferase reporter assays, plasmid production, plasmid transfection). YS, JR and FC were involved in NO and 3-NT determination. YS, CP and JR analyzed the data and performed bioinformatics analyses. BCC generated the NFAT5 KO cell lines. RF performed the NFAT5 rescue experiments and Western blot associated. The manuscript was written by CEI, FR, and FC with input from the other authors.

